# The what, how and why of trait-based analyses in ecology

**DOI:** 10.1101/2024.06.05.597559

**Authors:** Thomas Guillerme, Pedro Cardoso, Maria Wagner Jørgensen, Stefano Mammola, Thomas J. Matthews

## Abstract

Functional diversity is increasingly used alongside taxonomic diversity to describe populations and communities in ecology. Indeed, functional diversity metrics allow researchers to summarize complex occupancy patterns in space and/or time (what is changing?) that lead to changes in communities and/or populations (the process; how is it changing?) in response to some stressors (the mechanism; why is it changing?). However, as the diversity of functional diversity metrics and methods increases, it is often not directly clear which metric is more readily appropriate for which question. We studied the ability of different functional diversity metrics to recover patterns and signals from different processes linked to common assembly mechanisms (environmental filtering, competitive exclusion, equalizing fitness, and facilitation) in community ecology. Using both simulated data and an empirical dataset affected by more complex and nuanced mechanisms, we tested the effectiveness of different space occupancy metrics to recover the simulated or empirical changes. We show that different metrics perform better for different tasks, emphasizing the importance of not using a one-size-fits-all metric. Instead, researchers should carefully consider and test whether a particular metric will be effective in capturing a pattern of interest.

## 2 Introduction

In the last two decades, there has been a progressive expansion in ecology and evolution studies from taxonomic-oriented approaches, with species as the focal point, to functional approaches that place species-specific characteristics (traits) at the centre of analyses (Mammola et al., 2021; Palacio et al., 2022). Although numerous definitions exist (Dawson et al., 2021), we here consider a trait to be any observable characteristic (e.g. morphological, anatomical, ecological, physiological, behavioural, phenological, etc.) measured on individual organisms at any level, from genes to whole organisms. A trait-based approach has two advantages for answering core questions in ecology and evolution. First, it allows a deeper understanding of the mechanisms generating biodiversity patterns, by putting organisms’ traits at the centre of natural selection rather than the organisms themselves. Second, it allows comparisons across subdisciplines in biology, while facilitating the conceptualization of general principles broadly valid in space (e.g. unrelated species pools) and time (e.g. anatomical traits are comparable between palaeontology and ecology, which is not always the case with species; Luza et al. 2023). Already used routinely in palaeontology (Raup, 1961; Gould, 1991; Foote, 1995; Guillerme et al., 2020a), this new trait-focused ecology and evolution is unlocking the possibility to answer a broad range of questions in disciplines as diverse as community ecology (McGill et al., 2006), biogeography (Violle et al., 2014), conservation biology (Chichorro et al., 2022), micro- (Chapin III et al., 1993), macro-evolution (Guillerme et al., 2023) and applied fields (e.g. agronomy Martin and Isaac 2015). This is because trait-based approaches closely align with the general analytical framework proposed by Anand (1994) to answer three sequential questions: *“what?”* (describing the pattern), *“how?”* (describing the process) and *“why?”* (understanding the mechanism) (see Box 1).

### Box 1

Anand (1994) conceptualised a general analytical framework for answering scientific questions in ecology and evolution, based on three sequential questions: *“what?”* (describing the pattern), *“how?”* (describing the process) and *“why?”* (understanding the mechanism). This framework can be effectively applied to trait-based analyses:

*What?* Pattern description corresponds to the steps needed to collect and summarise data to answer a question of interest (i.e., “how?” and “why?” as defined below). Usually this consists in: i) collecting target traits at the focal level (e.g., genes, individual, population, species; Violle et al. 2007); ii) arranging these traits into some kind of trait space (Guillerme et al., 2020a; Mammola et al., 2021); and iii) using one or more unidimensional or multidimensional statistics to summarise properties of the trait space. For unidimensional spaces (i.e. distributions), these statistics can be as simple as the mean and the standard deviation (e.g. community weighted mean). For multidimensional spaces, these are usually named disparity metrics or indices (Guillerme et al., 2020a) or functional diversity metrics (Mammola et al., 2021), but all attempt to capture some pattern of interest in the trait space.

*How?* Process description can be seen as linking the pattern of interest (“What?”) to some dynamic element. This can be a punctual change. For example, change of the pattern under a certain condition, such as how traits X differ between two habitats (Martínez et al., 2021). But it can also be one or more continuous or ordinal changes, such as how the pattern X changes in space, time and/or along ecological gradients (Belmaker and Jetz, 2013; Jarzyna and Jetz, 2018; Lamanna et al., 2014; Bjorkman et al., 2018; McLean et al., 2021). The distinction here might seem trivial but it is significant: usually, the process designates the change of the pattern, not the change of the traits. Although the traits are what is really changing, researchers will usually analyse some emergent property of the trait aggregation as described by the change of statistics between two or more conditions.

*Why?* Mechanism description is then at the core of answering the biological question at hand. In most cases, this is what researchers are actually trying to understand. For example, one might be interested in understanding the effect of climate change on some trait (Boonman et al., 2022). In fact, most researchers work on understanding the causal link between variables (i.e. “why” are “what” and “how” linked). This can be very useful, for example, to provide predictions about the past or the future. Ultimately, studying the mechanisms (“why”) is often the reason why researchers get funded (or not).

Note that the distinction between what, how and why is not categorical in its nature and is often nuanced, with patterns and process or process and mechanisms sometimes being used interchangeably to describe the same questions.

While the first part of this framework (“what?”) has been studied extensively in the last decade, this was often undertaken outside of the “what, how and why” framework. Here we argue that the pattern description, although the basis of any future study, often follows some framework without a proper evaluation of its adequacy to answer the how and why. The choice of the tool or metric used to describe the pattern is crucial for allowing us to understand the process and the mechanism. Using an inappropriate metric for describing a pattern can lead to biased conclusions. For example, let us imagine one is interested in understanding how two populations compete with each other for a resource (the mechanism - “why?”) on two islands, one home to both populations and another one home to only one of the population (the process - “how?”). In such a scenario, measuring the occupied area (i.e. *km*^2^) of each population on each island (a pattern - “what?”) will not be the most appropriate way to understand the potential competition between these populations. In this simplistic example, the functional overlap between populations (“what and how”) might be more appropriate. See for example Carvalho and Cardoso 2020 for a discussion of how Darwin’s finches share resources depending on the existence or not of competition.

Through a simulation exercise, we analysed different patterns (what) across different processes (how) approximating different mechanisms (why) of interaction between organisms: equalisation, filtering, facilitation and competition (Figure 1). We also used an empirical dataset of Hawaiian bird traits and used pre-historical and historical extinction events on Hawaiian islands (process) resulting in a trait space modified by extinctions (the pattern) capturing the effects of species extinction on species trait distributions (the mechanism). We show that, with a fixed process and pattern, the choice of statistics used to describe the patterns (the disparity or functional diversity metrics) have a great impact on our ability to recover the process and understand the mechanism. We suggest caution when summarising patterns in observed data and propose tools for researchers to help understand how their pattern (often stemming from multidimensional data) can vary intuitively or not depending on the process and mechanism of interest.

**Figure 1:**
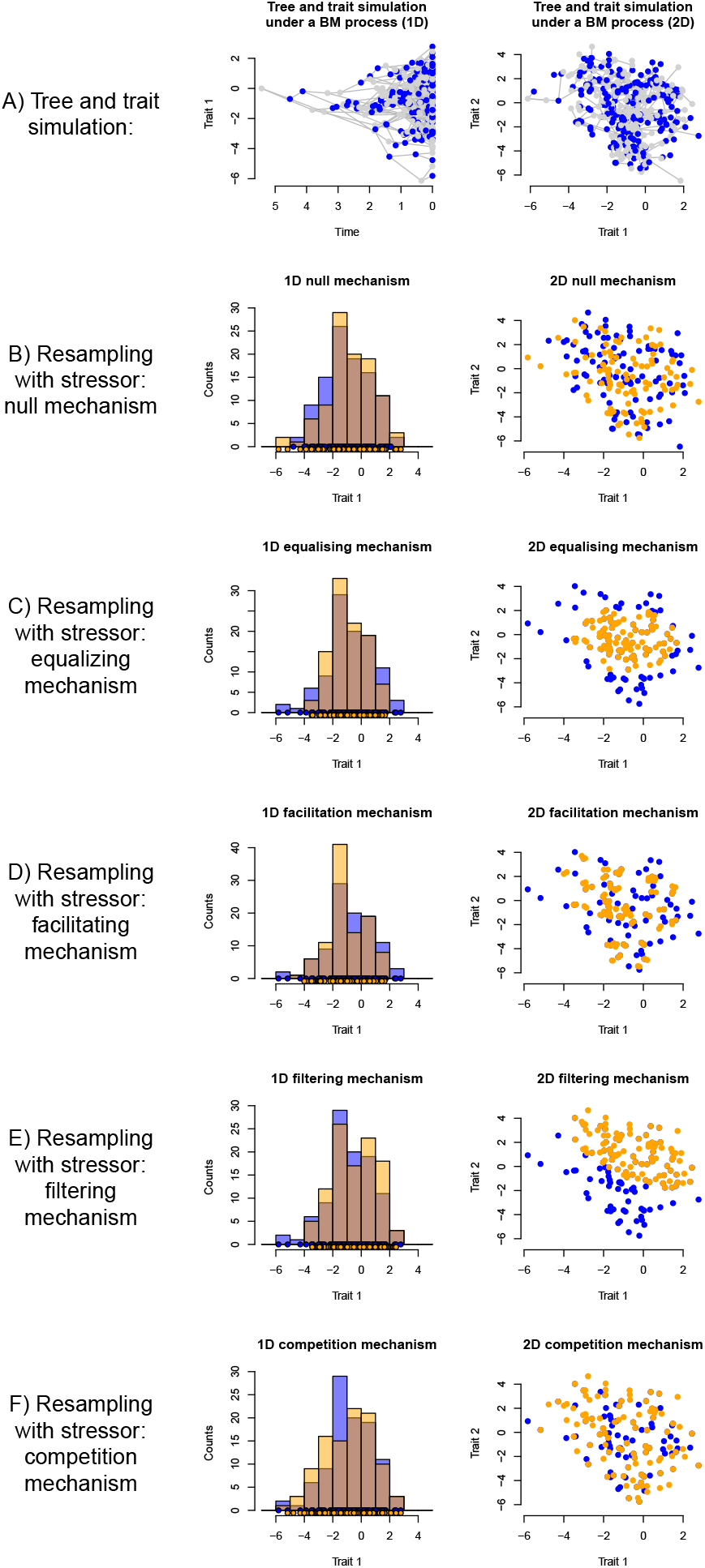
Illustration of the simulation protocol used in this paper. A) we simulated a pure birth-tree with a trait evolving under a Brownian Motion process until reaching 200 tips resulting a trait space of 2, 4, or 8 dimensions. We then applied different stressors to the resulting trait space using the following: B) **null mechanism**: randomly removing half of the species resulting in two groups of 100 tips (blue and orange - the brown colour in the histogram represents the overlap between both groups); C) **equalising mechanism**: removing species on the edge of the distribution; D) **facilitation mechanism**: removing species to reduce the distance between pairs of species (increasing local density; decreasing evenness); E) **filtering mechanism**: removing species on one extreme of the distribution; F) **competition mechanism**: proportionally removing species on the centre of the distribution (“flattening the curve”).

## 3 Methods

### 3.1 Simulating trait space patterns

First, we simulated a single Brownian Motion trait under a pure birth speciation model until reaching 200 elements (representing tips, individuals, species, OTUs, etc.) in R (R Core Team, 2024) using treats (Guillerme, 2024). This resulted in a neutral null model of trait evolution with no effect of competition, extinction, selection or other processes (*sensu* Bausman 2018). We refer to this as the “non-stressed trait space”.

#### 3.1.1 Applying stressors to the trait space

We then applied four different stressors to the non-stressed trait space to remove either 20%, 40%, 60% or 80% of the data (resulting in trait spaces with, respectively, 160, 120, 80 and 40 elements). In removing the data, we applied these five specific algorithms (Figure 1; Table 1; all algorithms, except “Evenness”, were previously described in Guillerme et al. (2020b)):

- Random removal: by randomly removing 20%, 40%, 60% or 80% of the data. This approximates our **null mechanism**.
- Decreasing size: by removing the required amount (20%, 40%, 60% or 80%) of data away from a distance (radius) *ρ* of the centre of the trait space. This approximates our **equalising mechanism**.
- Increasing density: by removing the required amount of pairs of points with a pair distance of at least *D* (i.e. removing the *n* pairs of points that are at least *D* distance away from each other). This approximates our **facilitation mechanism** (i.e. the points left are only ones that are close to at least another point in space).
- Shifting space: by removing the required amount of data from a distance (radius) *ρ* of the element with the maximum value. This approximates our **filtering mechanism**.
- Increasing evenness: by resampling the required amount of data but with skewed resampling probabilities per bandwidth (i.e. an estimated number of categories summarising the distribution, like in a histogram). Each bandwidth has an observed probability of sampling of *n* (proportional to the number of elements in
- that bandwidth) and is skewed to a probability of sampling of *n* × *m*^*p*^ where *m* > 0 for the low values of *n* and *m* < 0 for high values of *n* (respectively low and high density bandwidths) for flattening the curve (and the opposite for steepening it) and *p* is a factor increasing the flattening/steepening (here we used *p* = 3). This approximates our **competition mechanism**.

**Table 1:** Description of the stressors applied to the simulated data and the mechanism each approximated. Here we distinguish between the biological mechanism we are trying to simulate and the algorithmic one we used to simulate it. We also provide a more detailed description of the stressor. Note that in biological data, we don’t expect any of the mechanisms to act on communities alone (e.g. equalising fitness and facilitation can both act on the distribution of species traits). Nor do we expect their effects to be unidirectional (for example, equalising fitness can happen both by removing the edges or changing the position of a trait distribution). These mechanisms serve as a simplified description of reality for the narrative purpose of this paper.

| Approximated mechanism (biological) | Used mechanism (algorithm in `dispRity::reduce.space`) | Stressor description |
| --- | --- | --- |
| Null mechanism | Random removal ("random") | No overall change in species traits or community structure except for the reduction in the number of elements. |
| Equalising fitness | Size change ("size") | When species with trait combinations that are more extreme (i.e. species located away from the centre of the trait space) are disadvantaged due to some resource concentration gradient. |
| Facilitation | Density change ("density") | When species with closely shared trait combinations (i.e. located close together in the trait space) are more likely to survive the stressor. In other words, this simulates the idea that some species are more likely to be resilient if they are present in a community of species with shared traits rather than the opposite. |
| Environmental filtering | Position change ("position") | When an increase in species trait similarity happens through some strong abiotic constraints (Cornwell et al., 2006). For example, some trait combinations become more and more unlikely due to some environmental constraints (i.e. some regions of the trait space become unsuitable). |
| Competitive exclusion | Evenness change ("evenness") | When functionally similar species compete more intensively with one another than with functionally dissimilar species ("competitive exclusion principle"; Hardin 1960). At the extreme, it implies that only dissimilar individuals will coexist ("limiting similarity principle"; MacArthur and Levins 1967) |

**Table 2:**
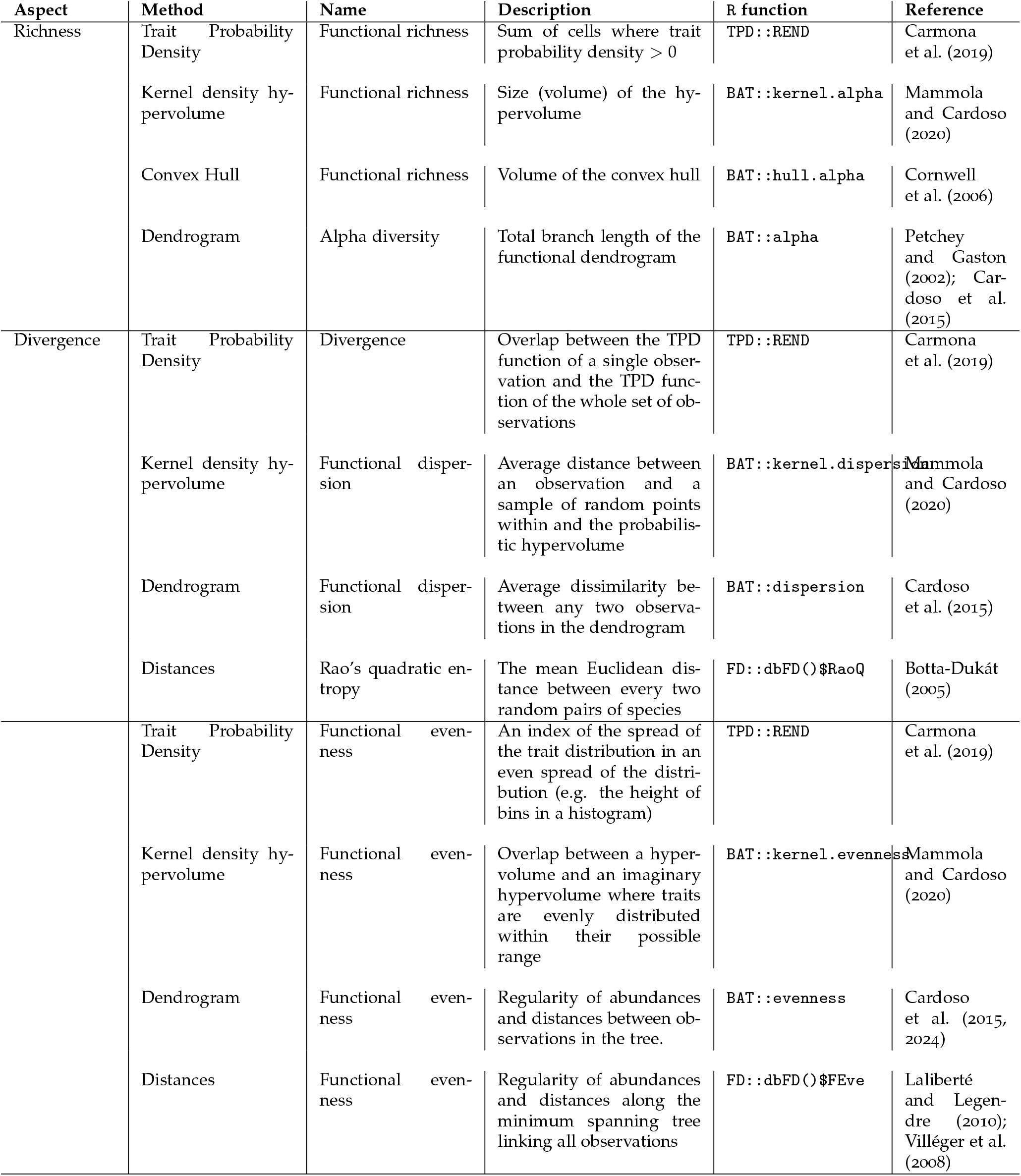
Functional diversity metrics tested in our simulations.

| Aspect | Method | Name | Description | R function | Reference |
| --- | --- | --- | --- | --- | --- |
| Richness | Trait Probability Density | Functional richness | Sum of cells where trait probability density $> 0$ | TPD::REND | Carmona et al. (2019) |
|  | Kernel density hypervolume | Functional richness | Size (volume) of the hypervolume | BAT::kernel.alpha | Mammola and Cardoso (2020) |
|  | Convex Hull | Functional richness | Volume of the convex hull | BAT::hull.alpha | Cornwell et al. (2006) |
|  | Dendrogram | Alpha diversity | Total branch length of the functional dendrogram | BAT::alpha | Petchey and Gaston (2002); Cardoso et al. (2015) |
| Divergence | Trait Probability Density | Divergence | Overlap between the TPD function of a single observation and the TPD function of the whole set of observations | TPD::REND | Carmona et al. (2019) |
|  | Kernel density hypervolume | Functional dispersion | Average distance between an observation and a sample of random points within and the probabilistic hypervolume | BAT::kernel.dispersion | Mammola and Cardoso (2020) |
|  | Dendrogram | Functional dispersion | Average dissimilarity between any two observations in the dendrogram | BAT::dispersion | Cardoso et al. (2015) |
| | Distances | Rao's quadratic entropy | The mean Euclidean distance between every two random pairs of species | FD::dbFD()\$RaoQ | Botta-Dukát (2005) |
|  | Trait Probability Density | Functional evenness | An index of the spread of the trait distribution in an even spread of the distribution (e.g. the height of bins in a histogram) | TPD::REND | Carmona et al. (2019) |
|  | Kernel density hypervolume | Functional evenness | Overlap between a hypervolume and an imaginary hypervolume where traits are evenly distributed within their possible range | BAT::kernel.evenness | Mammola and Cardoso (2020) |
|  | Dendrogram | Functional evenness | Regularity of abundances and distances between observations in the tree. | BAT::evenness | Cardoso et al. (2015, 2024) |
| | Distances | Functional evenness | Regularity of abundances and distances along the minimum spanning tree linking all observations | FD::dbFD()\$FEve | Laliberté and Legendre (2010); Villéger et al. (2008) |

For the position, size and density algorithms, we estimated the parameters *ρ* and *D* recursively with the `dispRity` package to obtain the required amount of data to be removed (`dispRity::reduce.space`, Guillerme 2018; Guillerme et al. 2020b). These steps resulted in 3420 simulated trait spaces (171 simulated spaces × 5 stressors × 4 amounts of data removed). We used 171 replicates per simulation because that was empirically the smallest number of simulations required to reach a variance between simulations lower than 1% across all metrics (i.e. any additional simulation added less that 1% extra variance). See Table 1 for a biological description of these stressors and Figure 1 for a visual description of them.

Note that in empirical data, depending on the distribution of the data, some mechanisms can lead to similar or dissimilar patterns. For example, if the data are normally distributed, the equalising mechanisms, by removing data on the edges of the distribution also increases the density of the trait space (because normally distributed data are denser in the centre of the distribution), similarly to the facilitation mechanism. However, if the data are distributed uniformly, this does not happen. Furthermore, in real-world scenarios, we do not expect these mechanisms to act in isolation of each other: multiple mechanisms may stress the observed data simultaneously, with cumulative or synergistic effects. However, this is not tackled here for both simplicity and to understand how they work in isolation.

We repeated each simulation pipeline (generate a trait space and apply the stressor) for uncorrelated 4 dimensional traits (Figure 2) and uncorrelated 2 and 8 dimensional traits (supplementary materials). We limited our simulations to a relatively small number of traits due to the constraints of some of the metrics used (e.g., `TPD::TPDsMean` is only implemented for up to 4 dimensions; Carmona et al. 2019) but also to avoid dealing with the curse of dimensionality (Bellman et al., 1957). This curse changes the properties of space occupancy in a non-linear way depending on the element’s distribution and the number of dimensions. For example, the volume of a trait space (or hypervolume when using more than 3 dimensions) typically tends to zero in a high number of dimensions. However, the rate at which it approaches zero is not linear and depends on the distribution of each element on every dimension. This makes it practically difficult to compare two randomly generated spaces with similar characteristics (e.g. for two spaces with 200 elements and 10 dimensions generated in the exact same way, one might have a hypervolume of nearly 0 and the other one of 10^5^). Note also that in our simulations, all dimensions have the same properties (i.e. same variance and distribution). This is often not the case in empirical cases (e.g. see our empirical example) where the dimensions have a decreased variance due to ordination techniques.

**Figure 2:**
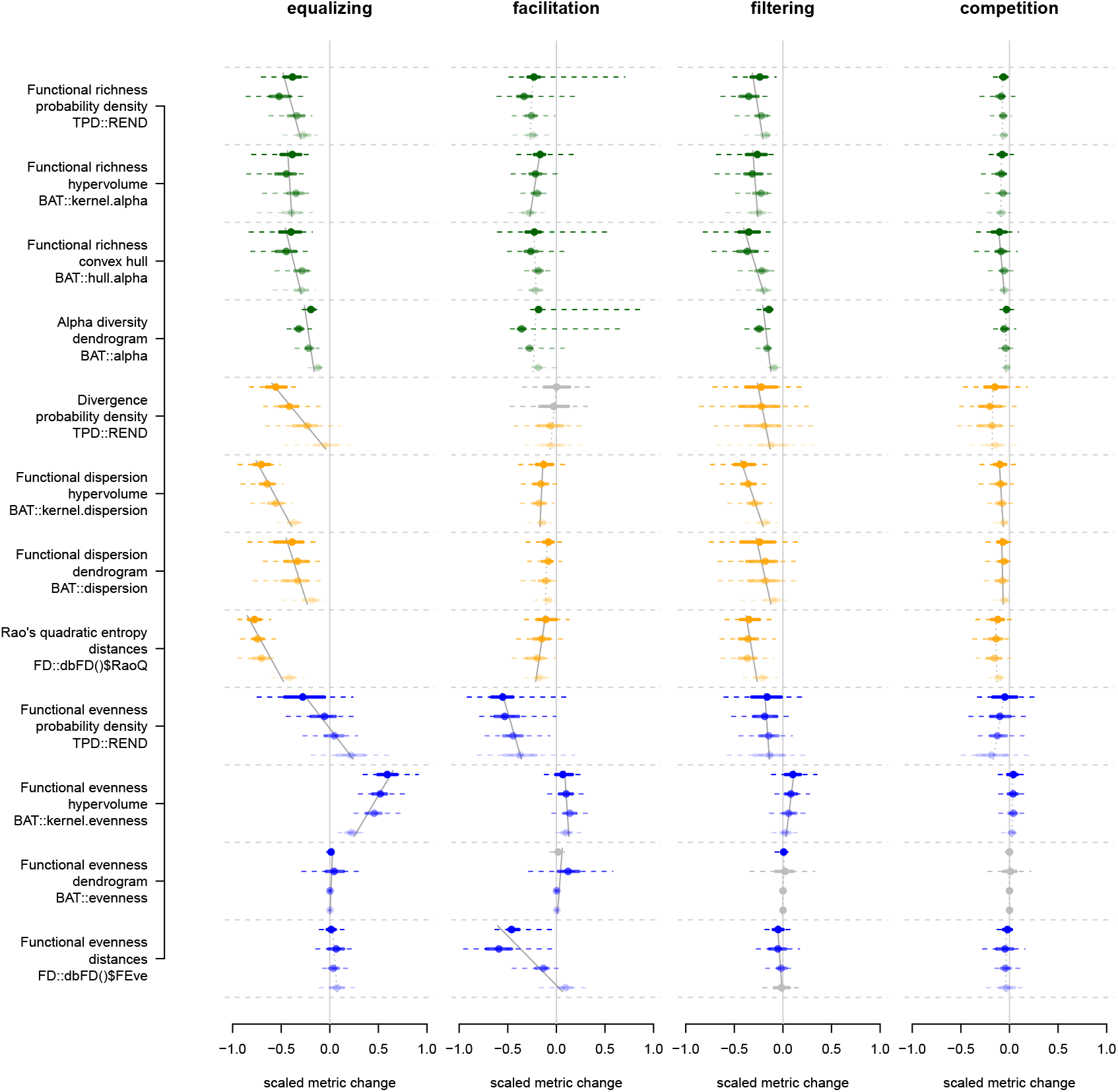
Simulation results: the y axes represent the different metrics tested (sorted by categories). The different columns represent the different stressors. The x-axes represent the metric values centred on the random changes and scaled by the maximum value for each metric between the four stressors. Negative and positive values signify a decrease/increase in the metric score. The dots represent the median metric value, the full line their 50% confidence interval (CI) and the dashed line their 95% CI. The colours are here to visually separate the metrics by categories (green = richness, yellow = divergence, blue = regularity); the colour gradient within each row corresponds to a removal of respectively 80%, 60%, 40% and 20% of the data (from top to bottom). The grey line plots represent distributions of metric scores not clearly distinguishable from the random metric scores (paired t-test *p* value > 0.05). Grey lines in the background across the distributions of different removal amounts represent the fitted linear model centred and scaled metric score *∼* amount of data removed and the value displayed is the adjusted *R*^2^ from each of these models. Dashed thin grey lines represent non-significant models (*p* value of slope or/and intercept > 0.05). Similar figures are available in the supplementary materials for 2 and 8 dimensions.

#### 3.1.2 Metrics for measuring trait space occupancy dissimilarity/disparity

We structured our simulations based on three aspects of diversity commonly captured by functional diversity metrics (Mammola et al., 2021): 1) Richness, encompassing metrics reflecting the sum of differences among observations; (4 metrics, equivalent to size metrics in Guillerme et al. 2020b); 2) Divergence, encompassing metrics reflecting the average differences among observations (4 metrics); and 3) Regularity, encompassing metrics reflecting how regular the differences among observations are (4 metrics, equivalent to density metrics in Guillerme et al. 2020b). Note that we focused here on three packages fully devoted to functional diversity analyses (BAT Cardoso et al. 2015 FD Laliberté and Legendre 2010 and TPD Carmona et al. 2019) in the R statistical environments (see Mammola et al. 2021).

#### 3.1.3 Scaling the results to be proportional to the random removals (null)

To ease interpretation of the results, we centred the results for the 12 metrics for the 4 non null stressors by the results for the null stressor and then scaled them. Effectively, for each metric and each single simulation, we subtracted the results under the null stressor to the ones under the other stressors and divided the resulting centred metrics by the maximum value for each metric between each simulation. This resulted in each metric being scaled between –1 and 1 where 0 corresponds to no change compared to the null and 1 and –1 the maximum positive and negative deviations relative to the null. In parallel, we also compared the distribution of the 171 scores for each metric with each reduction level for each non-null stressor to the null stressor (with no scaling or centering) using pairwise t-tests. The results of these tests are displayed in figure 2 (for the non-null stressor, grey distributions are not clearly distinguishable from the null stressor).

#### 3.1.4 Measuring the effect of the strength of the stressor

Finally, we measured the effect of the different levels of data removal (20%, 40%, 60% and 80%) by fitting a linear regression using the model centred and scaled metric score *∼* amount of data removed. For each of these models, we reported the model fit (adjusted *R*^2^) in the supplementary tables 1 to 3 and whether the slope was clearly distinguishable from 0 (Figure 2).

### 3.2 Empirical data

We applied all metrics to an empirical dataset of bird extinctions on six Hawaiian islands (Hawaii, Kauai, Lanai, Maui, Molokai, Oahu), an archipelago which has suffered large numbers of anthropogenic extinctions due to a range of extinction drivers (Walther and Hume, 2022). A prehistoric species list (avifauna known to be present prior to human colonisation of the islands), historic species list (avifauna known to be present at 1500 CE) and extant species list of the native community for each island was taken from Matthews et al. (2023) along with trait values for the extinct species (*Asio flammeus* was removed from the historic and extant species lists as its colonisation status is uncertain). We also added two marine species to the prehistoric species list; we compiled missing trait data for these two extinct species from Sayol et al. (2021). We extracted traits for extant species from the AVONET database (Tobias et al., 2022). We used 9 morphological traits associated with dietary/foraging preference and dispersal ability (Pigot et al., 2020; Sheard et al., 2020) to represent the functional diversity of the communities: mass; beak length (culmen); beak length (nares); beak width; beak depth; tarsus length; wing length; the length from the first secondary feather to the tip of the longest primary feather (Kipp’s distance); and tail length. Traits were log transformed and standardised to a mean of 0 and standard deviation of 1, prior to analyses.

The full trait dataset comprised 118 bird species, all native species known to have existed on these islands over the last 125,000 years. 55 species are known to have gone extinct prior to 1500 CE and 26 after 1500 CE. For our analyses, we focused on two time periods: (i) the avifauna present in 1500 CE (the “historic dataset”) and (ii) the current native avifauna (“extant dataset”). We represented the distinct reductions in species

richness preceding these two time periods as our two stressors, the first one representing all pre-1500 CE extinctions (i.e., 118 *™* 55 species; a species richness reduction of 46%) and the second all the extinctions that have occurred until the present (i.e., 118 *™* (55 + 26) species; a species richness reduction of 69%).

To build the community trait space, we undertook a PCA including all 118 species, which we then subset to calculate the trait space for the historic and extant datasets. We selected the first 5 axes to represent the trait spaces that explained at least 97.5% of the variance in each specific group (all, historic and extant species) We applied the same procedure as for the simulated data by simulating a null mechanism to compare to the observed ones by randomly removing 55 and 81 species for each stressor and scaling the results proportionally to this null mechanism (as described above).

## Results

The ability of different metrics to capture the different patterns (and thus approximate the processes) was highly variable. It ranged from metrics capturing no clear pattern (e.g. the Functional evenness based on the dendrogram method for competition) to metrics clearly capturing one specific pattern (e.g. Divergence based on probability density for the equalizing mechanism for a decrease in space occupancy). Some metrics even captured an opposite pattern to what was expected: for all metrics, we naively expected a relative decrease in trait space occupancy following different levels of species removals but some metrics captured an *increase* of trait space occupancy (e.g. the functional evenness based on hypervolume for the equalizing mechanism) or non-linear ones (e.g. Alpha diversity based on dendrogram for the equalizing mechanism). This raises an interesting point: the naive expectation that removing 80 to 20% of the data would result in a reduction of trait space occupancy metric score (negative values) is often true but by no means always true. Some cases resulted in an increase in metric score relative to random removals (positive values in Figure 2)!

For the empirical data, we see again a range of different possible interpretations depending on the metric describing the pattern. For example, all richness metrics (green; Figure 3) pick up a similar relative increase (i.e., relative to the null stressor) in richness for both the historic and extant datasets. However, the results are occasionally contrasting between the historical and extant datasets, with a respective decrease and increase with some divergence metrics (e.g. functional dispersion based on dendrograms or hypervolumes). Finally, some metrics pick up a decrease for both datasets (e.g. functional evenness based on hypervolumes - although the 95% confidence intervals overlapped with 0). Note that this apparent global increase in most metric values may appear counter intuitive given that 46% and 69% of species went extinct over the two time periods. This is due to the scaling of the metrics compared to random equivalent extinctions. That is, Figure 3 is not displaying absolute changes in trait space occupancy but rather changes in trait space occupancy relative to a crude neutral model where all species are equally likely to go extinct.

**Figure 3:**
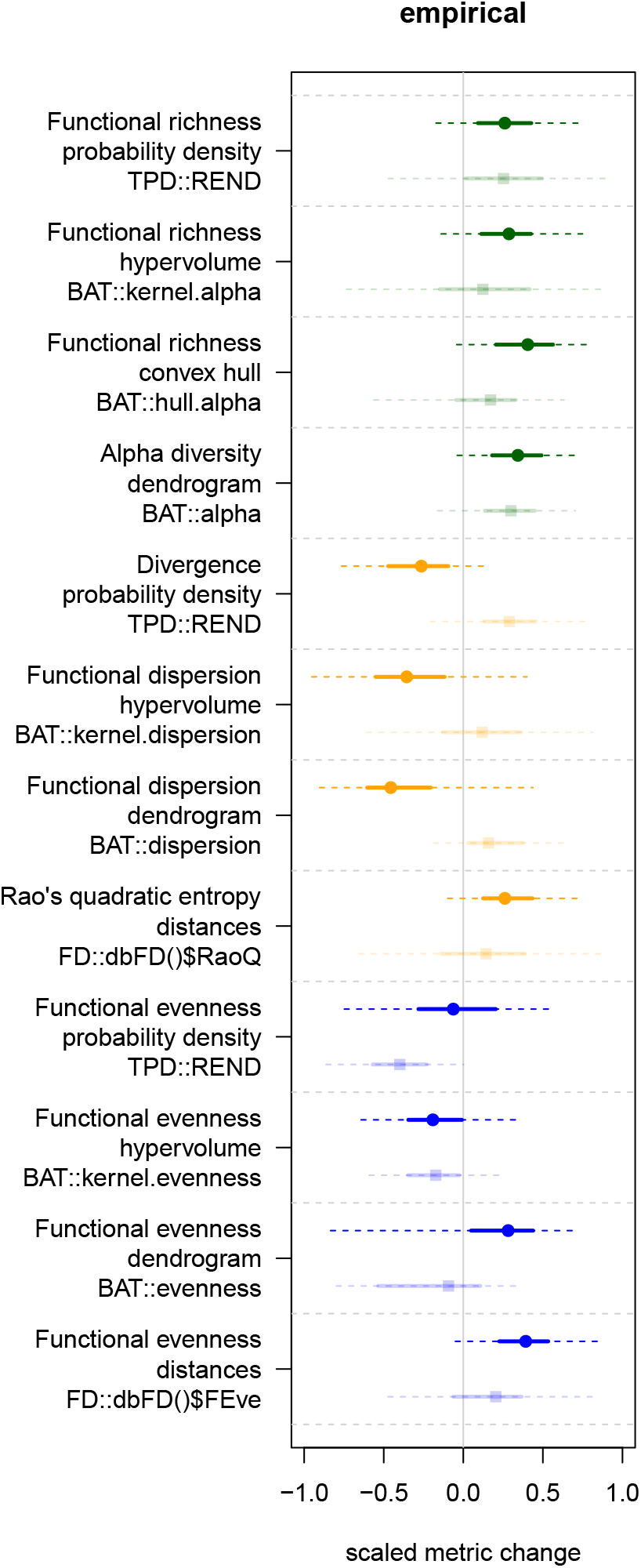
**Empirical results:** the metrics and the scaled changes are measured in the same way as in figure 2 (green = richness, yellow = divergence, blue = regularity). Plain circles represent the metrics for the historic dataset (46% removal) and fainted squares for the extant dataset (69% removal).

More broadly, as can be seen from this empirical case, patterns are likely to be much less clear than with simulated data due to multiple stressors acting simultaneously. For instance, extensive hunting is likely to present equalising and/or filtering pressure due to the selective hunting of larger bodied species. In contrast, avian malaria, which is known to be an important extinction pressure of native Hawaiian birds (Samuel et al., 2011), could show signals of a null mechanism since susceptibility to the disease is not trait-dependent (at least not in relation to the traits we have considered here), but is related to genetics (it mainly affects passerines, but there is considerable variation in susceptibility within passerines) and distributional range (Atkinson, 2023).

## Discussion

We tested 12 trait space occupancy metrics on simulated and empirical datasets to assess how each metric captures patterns of trait space changes based on stressors approximating ecological and evolutionary mechanisms. Our results show that, unexpectedly, different metrics capture different patterns (what) leading to inferring different processes (how) that can be proposed to explain different mechanisms (why). Therefore, the choice of trait space occupancy metric is essential to accurately describe a pattern of interest.

Our results may be influenced by our choice of space occupancy metrics - specifically, we focused on metrics available in three statistical packages in R, the most common statistical language currently used in ecology (Lai et al., 2019), but many others could have been used (e.g. Guillerme et al. 2020b). Also, our results are likely affected by the use of simplified algorithms designed to approximate very complex ecological and evolutionary mechanisms by simply removing a percentage of elements in trait spaces in a non-random manner. Note also that the simulated trait spaces here did not always share common characteristics with empirical trait spaces. The simulated trait spaces in 2, 4 or 8 dimensions had the same variance on all dimensions (i.e. we used the same trait simulation process for all dimensions) whereas in empirical trait spaces, commonly generated using ordination techniques (e.g. PCA or PCO), the dimensions have by definition a decreasing variance.

Furthermore, we only presented results based on a relatively small number of dimensions (up to 8), while it is not uncommon to have a much greater number of dimensions, especially in paleontology (e.g. in Van Den Ende et al. 2023). In a higher number of di-mensions (usually >10 but this is highly variable depending on the system under study), metrics results are harder to scale linearly. This sometimes makes metric interpretation more chaotic Bellman et al. (1957) in the sense that there is no direct relationship between the number of observations and dimensions that can be used to extrapolate results in higher dimensional datasets. In fact, the relationship between observations has an important effect on how our results scale with higher dimensionality (e.g. the volume of an evenly occupied space scales differently in high dimensions compared to the same space but with normally distributed observations). Therefore we advise workers to test assumptions specifically about their trait space and “play” around with it before choosing their functional diversity/disparity framework and metric of interest.

Another cautionary note could be made about our choice of scaling the metrics relative to a random trait space reduction (with the same percentage of reduction). This choice allowed us to simulate a null model designed to test whether the mechanisms of interest were causing the observed change in pattern (Bausman, 2018). For example, if removing 80% of the edges of a distribution (equalising mechanism) was distinguishable or not from removing 80% of a distribution randomly (null mechanism). Some metrics (e.g. Functional evenness using the dendrogram method) were not able to distinguish between the null mechanism and the mechanism of interest (here the mechanism simulating competition). In other words, our statistical question was “Does metric X distinguish between removing N% of data in a biassed way (the mechanism) and removing N% of data randomly (the null)?”. This definition of null hypothesis is also appropriate for the empirical data if the question is the same. However, it is very likely that workers will ask a more exciting or intriguing question based on these data. For example, an interesting one could be “Are the extinctions uniform across all birds in Hawaii?”. This legitimate question would thus require first a trait space (the pattern - what) and maybe some contrasting groups of interest like extinctions through time (the process - how) to answer the question (the mechanism - why). Furthermore, the scaling of the metrics paired with the levels of removal (80%, 60%, 40% and 20%) creates an expected artifact similarity when comparing the random removals against the other ones: when less data is available overall, the metrics are more likely to be similar (e.g. if removing 99% of the data, we expect the random and removal metric score to be nearly identical to the non-random ones).

The empirical results are also intriguing, as it indicates that, at least for certain metrics, the loss of Hawaiian bird functional diversity is less than expected than if extinctions were random with respect to traits (Figure 3). This contrasts with studies that have shown that species with certain traits (e.g., larger species) are more likely to have gone extinct and thus that functional diversity loss is larger than through random extinctions (e.g. Sayol et al. 2021; Matthews et al. 2022). This could partly be due to the proportion of passerine extinctions in Hawaii. Although most global bird species are (or were, if accounting for anthropogenic extinctions) passerines (*∼*58%), the majority of global known anthropogenic extinctions (*∼*75%) have been non-passerines, which tend to be larger and possess more unique morphological traits (thus resulting in relatively large reductions in functional diversity). However, in Hawaii, where the same proportion of known species were passerines (*∼*58%), the opposite is true: the majority of extinctions (*∼*60%) have been of passerine species. It is important to note that we are not implying that there existed little morphological variation within extinct Hawaiian passerines - indeed, many of these extinctions involved the honeycreepers, a famous adaptive radiation involving substantial evolution of beak morphology (Walther and Hume, 2022) - but simply that this morphological variation is less than that observed across all birds. But these results, given that the proportion of known Hawaiian species that are / were passerines roughly matches the proportion of extinct species that were passerines, also indicate that these extinct species were more functionally similar than expected, thus relatively *increasing* trait diversity (i.e. relative to random extinctions). In addition, it is worth stressing that these analyses included marine species (which constitute less than 4% of known extinctions in Hawaii, while making up 11% of the prehistoric assemblage) and different results may have been found if focusing exclusively on terrestrial birds.

Following our results and these thoughts, we encourage workers in ecology and evolution to test whether their metrics are capable of capturing a pattern of interest before discussing the process and/or mechanism inferred by it. We encourage workers to test their metrics using simplified example datasets to see if they have the ability to capture at least crude changes as exemplified in this work (e.g. using simulations Guillerme 2024 or the `moms` interactive package Guillerme et al. 2020b). Importantly, it should always be borne in mind that empirical mechanisms ought of course to be more complex in real world scenarios. For example, evolutionary mechanisms can vary through time or per clades (and so patterns are often the results of multiple processes); or ecological mechanisms are often intertwined and work together to generate a pattern (e.g. facilitation + competition; Danet et al. 2024), or counteract each others by operating on the trait space in opposite directions (e.g., competition + filtering; Mammola et al. 2024)

## Conclusion

Different ecological and evolutionary processes or mechanisms do not always results in different patterns. Our results based on simulations and empirical data suggest that the same data can be interpreted differently depending on the choice of trait space occupancy metric (*aka* disparity, dissimilarity or functional diversity metrics). Different metrics are designed to capture different aspects (Guillerme et al., 2020b; Mammola et al., 2021) but also perform better or worse at their designed task depending on the data, the mechanism or/and process of interest. Using space occupancy metrics for describing functional diversity is tricky. This is because the metrics have properties that can be counter intuitive based on the data at hand. And also because functions of ecosystems or organisms and niches within it are hard to capture/understand/define. This means that measuring functional diversity is based on both the definition of the metric and the function of interest and thus cannot be served by a one-fits-all metric. In other words, it

is important to choose a disparity/dissimilarity/functional diversity metric depending on the question and the data at hand rather than as a default option or solely based on a previous inspiring publication.

## 7 Repeatability and reproducibility

The simulations, results figures and tables are entirely reproducible via https://github.com/TGuillerme/eco_metrics_simulations. An additional vignette on how to choose metrics is available at https://github.com/TGuillerme/eco_metrics_simulations/blob/master/Analysis/Choosing_metrics_vignette.Rmd.

## 8 Acknowledgements

Thanks to Andrew Beckerman, Natalie Cooper, Alain Danet, Thomas Johnson and Gavin Thomas for comments on early versions of this manuscripts. TG was funded by the UKRI-NERC grant NE/X016781/1. SM was supported by NBFC, funded by the Italian Ministry of University and Research, PNRR, Missione 4, Componente 2, “Dalla ricerca all’impresa”, Investimento 1.4, Project CN00000033. MWJ was supported by NERC CENTA2 grant NE/S007350/1 and University of Birmingham.

## Notes

https://github.com/TGuillerme/eco_metrics_simulations/

### Competing Interest Statement

The authors have declared no competing interest.

