## Supplementary materials for "The what, how and why of trait-based analyses in ecology"

Version dated: June 5, 2024

### 1 Supplementary materials

#### 1.1 Supplementary results

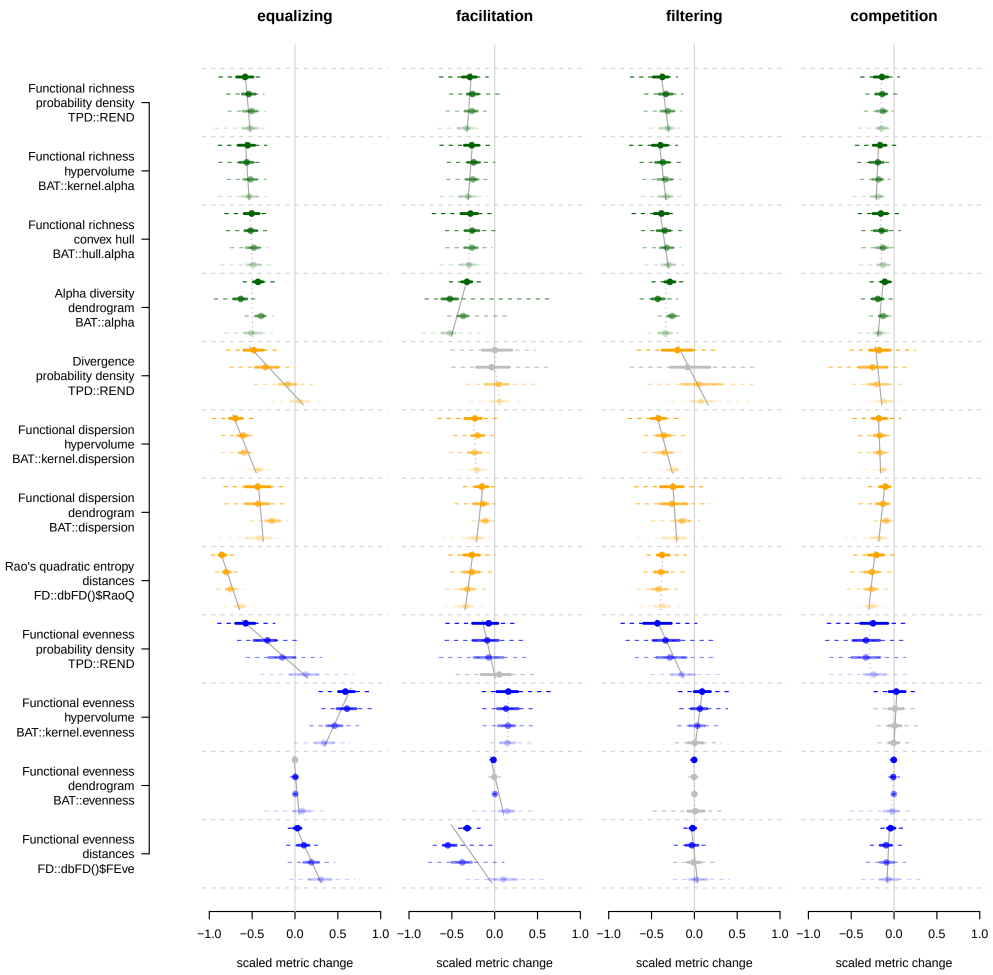


Figure 1: **Simulation results for 2 dimensions:** the y axes represent the different metrics tested (sorted by categories). The different columns represent the different stressors. The x-axes represent the metric values centred on the random changes and scaled by the maximum value for each metric between the four stressors. Negative and positive values signify a decrease/increase in the metric score. The dots represent the median metric value, the full line their 50% confidence interval (CI) and the dashed line their 95% CI. The colours are here to visually separate the metrics by categories (blue = regularity, yellow = divergence, green = richness); the colour gradient within each row corresponds to a removal of respectively 80%, 60%, 40% and 20% of the data (from top to bottom). The grey line plots represent distributions of metric scores not clearly distinguishable from the random metric scores (paired t-test p value $> 0.05$). Grey lines in the background across the distributions of different removal amounts represent the fitted linear model centred and scaled metric score $\sim$ amount of data removed and the value displayed is the adjusted $R^2$ from each of these models. Dashed thin grey lines represent non-significant models (p value of slope or/and intercept $> 0.05$).

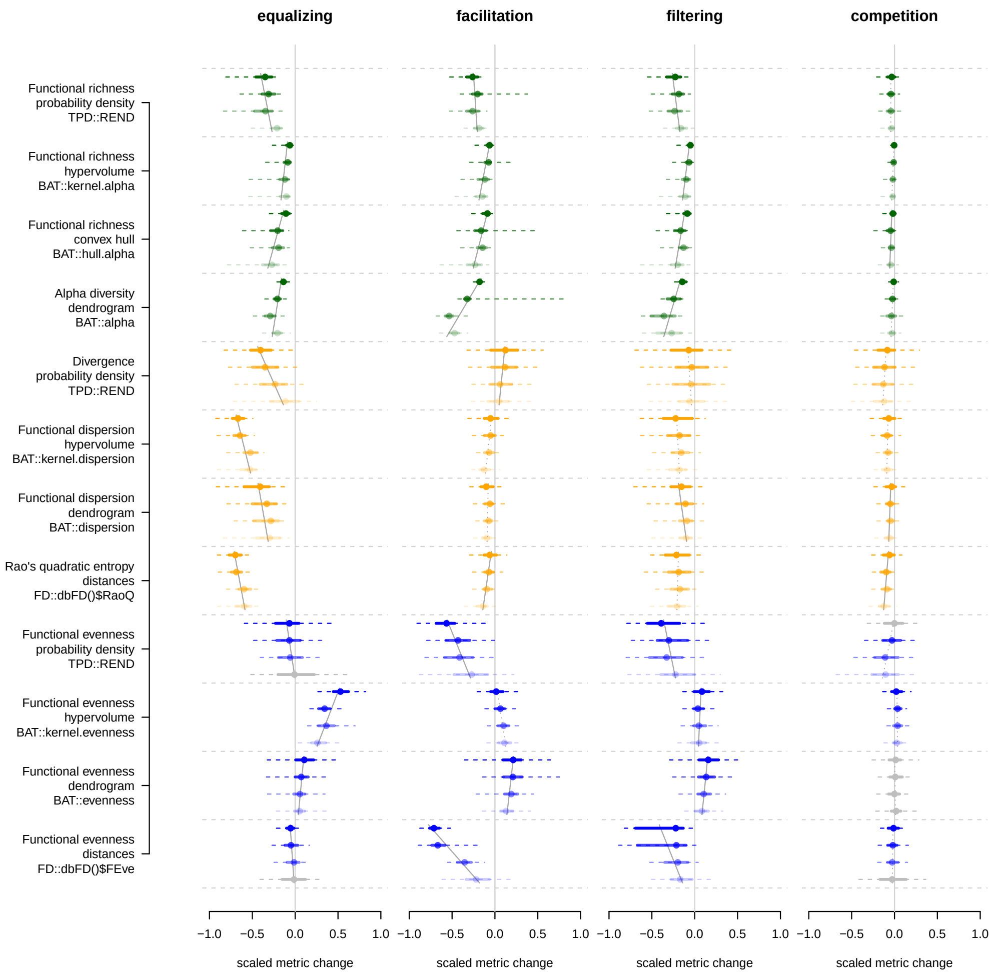


Figure 2: **Simulation results for 8 dimensions:** the y axes represent the different metrics tested (sorted by categories). The different columns represent the different stressors. The x-axes represent the metric values centred on the random changes and scaled by the maximum value for each metric between the four stressors. Negative and positive values signify a decrease/increase in the metric score. The dots represent the median metric value, the full line their 50% confidence interval (CI) and the dashed line their 95% CI. The colours are here to visually separate the metrics by categories (blue = regularity, yellow = divergence, green = richness); the colour gradient within each row corresponds to a removal of respectively 80%, 60%, 40% and 20% of the data (from top to bottom). The grey line plots represent distributions of metric scores not clearly distinguishable from the random metric scores (paired t-test p value $> 0.05$). Grey lines in the background across the distributions of different removal amounts represent the fitted linear model centred and scaled metric score $\sim$ amount of data removed and the value displayed is the adjusted $R^2$ from each of these models. Dashed thin grey lines represent non-significant models (p value of slope or/and intercept $> 0.05$).

Table 1: Results of the model scaled metric $\sim$ removal level per stressor (2D)

| | equalizing slope | equalizing adj.$R^2$ | facilitation slope | facilitation adj.$R^2$ | filtering slope | filtering adj.$R^2$ | competition slope | competition adj.$R^2$ |
|---|---|---|---|---|---|---|---|---|
| Functional evenness distances `FD::dbFD()$FEve` | 0.017*** | 0.02 | -0.015** | 0.012 | 0.026*** | 0.057 | -0.001 | -0.001 |
| Functional evenness dendrogram `BAT::evenness` | 0.014** | 0.011 | -0.016** | 0.014 | 0.024*** | 0.038 | -0.009* | 0.007 |
| Functional evenness hypervolume `BAT::kernel.evenness` | 0.007 | 0.002 | -0.006 | 0 | 0.03*** | 0.069 | 0.004 | 0 |
| Functional evenness probability density `TPD::REND` | 0.002 | -0.001 | -0.055*** | 0.07 | 0.001 | -0.001 | -0.02*** | 0.048 |
| Rao's quadratic entropy distances `FD::dbFD()$RaoQ` | 0.185*** | 0.578 | 0.022** | 0.009 | 0.102*** | 0.117 | 0.022** | 0.013 |
| Functional dispersion dendrogram `BAT::dispersion` | 0.08*** | 0.396 | 0.009. | 0.003 | 0.054*** | 0.195 | 0.008* | 0.005 |
| Functional dispersion hypervolume `BAT::kernel.dispersion` | 0.018** | 0.009 | -0.023*** | 0.04 | 0.014* | 0.004 | -0.022*** | 0.048 |
| Divergence probability density `TPD::REND` | 0.066*** | 0.436 | -0.028*** | 0.046 | -0.009. | 0.004 | -0.022*** | 0.038 |
| Alpha diversity dendrogram `BAT::alpha` | 0.227*** | 0.572 | 0.044*** | 0.037 | 0.091*** | 0.155 | 0.002 | -0.001 |
| Functional richness convex hull `BAT::hull.alpha` | -0.09*** | 0.293 | -0.002 | -0.001 | -0.027*** | 0.043 | -0.011* | 0.008 |
| Functional richness hypervolume `BAT::kernel.alpha` | 0.018*** | 0.049 | 0.046*** | 0.179 | 0.004 | 0 | -0.012*** | 0.02 |
| Functional richness probability density `TPD::REND` | 0.09*** | 0.356 | 0.14*** | 0.28 | 0.022*** | 0.037 | -0.009* | 0.005 |

Table 2: Results of the model scaled metric $\sim$ removal level per stressor (4D)

| | equalizing slope | equalizing adj.$R^2$ | facilitation slope | facilitation adj.$R^2$ | filtering slope | filtering adj.$R^2$ | competition slope | competition adj.$R^2$ |
|---|---|---|---|---|---|---|---|---|
| Functional evenness distances `FD::dbFD()$FEve` | 0.054*** | 0.143 | -0.009 | 0.002 | 0.032*** | 0.076 | 0.002 | 0 |
| Functional evenness dendrogram `BAT::evenness` | 0.013* | 0.007 | -0.032*** | 0.092 | 0.015** | 0.012 | -0.002 | 0 |
| Functional evenness hypervolume `BAT::kernel.evenness` | 0.049*** | 0.122 | 0.008 | 0.002 | 0.061*** | 0.194 | 0.018*** | 0.038 |
| Functional evenness probability `TPD::REND` | 0.03*** | 0.135 | -0.01 | 0.002 | 0.024*** | 0.09 | 0.001 | -0.001 |
| Rao's quadratic entropy distances `FD::dbFD()$RaoQ` | 0.164*** | 0.539 | -0.02** | 0.012 | 0.039*** | 0.023 | -0.002 | -0.001 |
| Functional dispersion dendrogram `BAT::dispersion` | 0.108*** | 0.535 | -0.009** | 0.009 | 0.067*** | 0.258 | 0.012*** | 0.025 |
| Functional dispersion hypervolume `BAT::kernel.dispersion` | 0.064*** | 0.145 | -0.004 | 0.002 | 0.045*** | 0.05 | 0.005* | 0.004 |
| Divergence probability density `TPD::REND` | 0.11*** | 0.52 | -0.03*** | 0.071 | 0.034*** | 0.065 | 0.004 | 0.001 |
| Alpha diversity dendrogram `BAT::alpha` | 0.153*** | 0.393 | 0.059*** | 0.083 | 0.014* | 0.005 | -0.035*** | 0.048 |
| Functional richness convex hull `BAT::hull.alpha` | -0.118*** | 0.517 | 0.013*** | 0.018 | -0.025*** | 0.06 | -0.002 | 0 |
| Functional richness hypervolume `BAT::kernel.alpha` | -0.008** | 0.009 | -0.015*** | 0.016 | -0.006 | 0.002 | -0.001 | -0.001 |
| Functional richness probability density `TPD::REND` | 0.016*** | 0.037 | 0.196*** | 0.535 | 0.014*** | 0.02 | -0.001 | -0.001 |

Table 3: Results of the model scaled metric $\sim$ removal level per stressor (8D)

| | equalizing slope | equalizing adj.$R^2$ | facilitation slope | facilitation adj.$R^2$ | filtering slope | filtering adj.$R^2$ | competition slope | competition adj.$R^2$ |
|---|---|---|---|---|---|---|---|---|
| Functional evenness distances `FD::dbFD()$FEve` | 0.039*** | 0.071 | 0.013** | 0.009 | 0.026*** | 0.051 | 0 | -0.001 |
| Functional evenness dendrogram `BAT::evenness` | -0.023*** | 0.041 | -0.038*** | 0.122 | -0.024*** | 0.102 | -0.008*** | 0.056 |
| Functional evenness hypervolume `BAT::kernel.evenness` | -0.055*** | 0.159 | -0.051*** | 0.121 | -0.034*** | 0.093 | -0.007*** | 0.015 |
| Functional evenness probability `TPD::REND` | -0.032*** | 0.134 | -0.118*** | 0.345 | -0.059*** | 0.185 | -0.008*** | 0.03 |
| Rao's quadratic entropy distances `FD::dbFD()$RaoQ` | 0.088*** | 0.157 | -0.02** | 0.01 | 0.013 | 0.002 | -0.012. | 0.004 |
| Functional dispersion dendrogram `BAT::dispersion` | 0.05*** | 0.166 | -0.022*** | 0.055 | 0.01. | 0.003 | -0.005 | 0.002 |
| Functional dispersion hypervolume `BAT::kernel.dispersion` | 0.035*** | 0.035 | -0.004 | 0.001 | 0.028*** | 0.029 | -0.007** | 0.009 |
| Divergence probability density `TPD::REND` | 0.037*** | 0.121 | -0.029*** | 0.101 | -0.005 | 0 | -0.017*** | 0.041 |
| Alpha diversity dendrogram `BAT::alpha` | 0.03*** | 0.019 | 0.08*** | 0.131 | 0.041*** | 0.028 | -0.041*** | 0.05 |
| Functional richness convex hull `BAT::hull.alpha` | -0.077*** | 0.265 | 0.031*** | 0.09 | -0.009* | 0.006 | 0.002 | -0.001 |
| Functional richness hypervolume `BAT::kernel.alpha` | -0.019** | 0.013 | -0.023*** | 0.017 | -0.02*** | 0.018 | 0.002 | -0.001 |
| Functional richness probability density `TPD::REND` | 0.014** | 0.012 | 0.175*** | 0.574 | 0.08*** | 0.112 | -0.002 | -0.001 |

### References
